# Early resource scarcity causes cortical astrocyte enlargement and sex-specific changes in the orbitofrontal cortex transcriptome in adult rats

**DOI:** 10.1101/2023.07.01.547315

**Authors:** Claire Deckers, Reza Karbalaei, Nylah A. Miles, Eden V. Harder, Emily Witt, Erin P. Harris, Kathryn Reissner, Mathieu E. Wimmer, Debra A. Bangasser

**Author notes:** Correspondence: Debra Bangasser, PhD, Georgia State University 100 Piedmont Ave., SE Atlanta, GA 30303.

## Abstract

Astrocyte morphology affects function, including the regulation of glutamatergic signaling. This morphology changes dynamically in response to the environment. However, how early life manipulations alter adult cortical astrocyte morphology is underexplored. Our lab uses brief postnatal resource scarcity, the limited bedding and nesting (LBN) manipulation, in rats. We previously found that LBN promotes later resilience to adult addiction-related behaviors, reducing impulsivity, risky decision-making, and morphine self-administration. These behaviors rely on glutamatergic transmission in the medial orbitofrontal (mOFC) and medial prefrontal (mPFC) cortex. Here we tested whether LBN changed astrocyte morphology in the mOFC and mPFC of adult rats using a novel viral approach that, unlike traditional markers, fully labels astrocytes. Prior exposure to LBN causes an increase in the surface area and volume of astrocytes in the mOFC and mPFC of adult males and females relative to control-raised rats. We next used bulk RNA sequencing of OFC tissue to assess transcriptional changes that could increase astrocyte size in LBN rats. LBN caused mainly sex-specific changes in differentially expressed genes. However, *Park7*, which encodes for the protein DJ-1 that alters astrocyte morphology, was increased by LBN across sex. Pathway analysis revealed that OFC glutamatergic signaling is altered by LBN in males and females, but the gene changes in that pathway differed across sex. This may represent a convergent sex difference where glutamatergic signaling, which affects astrocyte morphology, is altered by LBN via sex-specific mechanisms. Collectively, these studies highlight that astrocytes may be an important cell type that mediates the effect of early resource scarcity on adult brain function.

## 1.0 Introduction

Astrocytes are prevalent glial cells in the brain that have important homeostatic functions, regulating synaptic transmission, synapse formation and remodeling, and metabolism (Sofroniew and Vinters 2010, Clarke and Barres 2013, Verkhratsky and Parpura 2016, Kim, Healey et al. 2018). One key feature that influences the function of astrocytes and the neurons they regulate is their morphology. Astrocytes regularly adjust their morphology to support dynamic brain functions, such as the neurovascular coupling and synaptic remodeling that occurs during sleep or lactation (Bellesi, de Vivo et al. 2015, Li, Li et al. 2021). Increasing astrocyte size can be beneficial, providing trophic support to neurons. However, reactive astrocytes with a hypertrophic morphology can disrupt domain architecture and, in some cases, form a glial scar that prevents contact between the damaged area and adjacent tissue (Sofroniew and Vinters 2010, Verkhratsky and Parpura 2016). In contrast, atrophic astrocytes that are smaller and less arborized than typical astrocytes can reduce the tone of signaling molecules, homeostatic capabilities, and network connectivity (Pekny, Pekna et al. 2016, Verkhratsky and Parpura 2016, Kim, Healey et al. 2018). Given the importance of astrocytic morphology to overall brain function, it is crucial to understand how the changes in the environment impact the shape of astrocytes.

Stressful events and stress-related disorders can alter cortical astrocytes (for review see (Wang, Jie et al. 2017, Kim, Healey et al. 2018, Abbink, van Deijk et al. 2019, Murphy-Royal, Gordon et al. 2019). For example, in postmortem tissue from people (particularly young people) with major depression (MDD), there is a reduction in the astrocyte-specific marker, glial-fibrillary acid protein (GFAP), in regions including the prefrontal cortex (PFC), dorsolateral (dl) PFC, anterior cingulate cortex (ACC), and orbitofrontal cortex (OFC) (Miguel-Hidalgo, Baucom et al. 2000, Webster, Knable et al. 2001, Si, Miguel-Hidalgo et al. 2004, Miguel-Hidalgo, Waltzer et al. 2010, Gittins and Harrison 2011). Morphological analyses of astrocytes in postmortem tissue are limited. However, one study reported reduced glial size in the gray matter of caudal OFC and dlPFC in the MDD versus control groups (Rajkowska, Miguel-Hidalgo et al. 1999), indicating that MDD causes a reduction in cortical astrocyte number and morphology.

Rodent studies exposing adult animals to chronic stressors (e.g., restraint and chronic mild stress) to mimic some aspects of depression-related outcomes similarly find that chronic stress reduces GFAP in the frontal cortex (Banasr, Chowdhury et al. 2010, Sántha, Veszelka et al. 2016, Shilpa, Bhagya et al. 2017). In macaque monkeys that exhibit anxiety-related self-injurious behaviors, astrocyte morphology (e.g., arbor length and bifurcations) is reduced in the frontal cortex (Lee, Chiu et al. 2013). In male rats, chronic restraint stress in adulthood causes atrophy of astrocytes in the PFC but does not alter astrocyte number suggesting that the shape of astrocytes may be preferentially regulated by stress (Tynan, Beynon et al. 2013, Bollinger, Salinas et al. 2019). In contrast in female rats, chronic restraint stress causes astrocyte hypertrophy in the PFC, an effect linked to ovarian hormones (Bollinger, Salinas et al. 2019), underscoring the need to compare environmental effects on astrocytes across sex.

Much of the work on stress and cortical astrocytes has focused on chronic stress in adult animals. However, early life stress is a risk factor for depression and many other psychiatric disorders including substance use disorder (SUD) (Baskin-Sommers and Foti 2015, Reiss, Meyrose et al. 2019, LeMoult, Humphreys et al. 2020, al’Absi, Ginty et al. 2021). In rats, maternal separation reduces GFAP in the medial (m) PFC a day after the final separation and this effect appears to last into adulthood (Leventopoulos, Rüedi-Bettschen et al. 2007, Musholt, Cirillo et al. 2009, Banqueri, Méndez et al. 2019). Few studies have assessed whether early life stress alters astrocyte morphology, however acute family separation stress reduces the GFAP immunoreactive processes in the mPFC of juvenile *Octodon degus* (Braun, Antemano et al. 2009). These studies do suggest that early life stress has the potential to cause lasting changes in cortical astrocytes, but there are few, and focus on separation stress. Thus, it is difficult to determine if this work translates to other types of early life stress.

Our laboratory uses the limited bedding and nesting (LBN) model of early adversity, where dams and pups are put in a low resource environment from pups’ postnatal day (PND) 2-9. This manipulation fragments maternal care and increases dams’ pup-directed behavior at the expense of their self-care, which is likely reflective of a hyperarousal phenotype (Ivy, Brunson et al. 2008, Walker, Bath et al. 2017, Eck, Ardekani et al. 2019, Gallo, Shleifer et al. 2019, Shupe and Clinton 2021, Wendel, Short et al. 2021). In our hands, we have found the lasting effects of LBN on offspring to be largely protective, likely due to the brief nature of the stressor and the intense, albeit altered, maternal care. Specifically, we have determined that LBN reduces impulsivity, as measured with delayed discounting, and morphine self-administration in adult male rats, and causes adult males and females to be more risk averse in a probability discounting task (Ordoñes Sanchez 2021, Ordoñes Sanchez, Bavley et al. 2021). Two key brain regions engaged by these behaviors are mPFC and the OFC (Bechara, Tranel et al. 2000, Mobini, Body et al. 2002, Pattij and Vanderschuren 2008, St. Onge and Floresco 2010, Mar, Walker et al. 2011, Weidacker, Johnston et al. 2020), areas where other types of stressors are known to affect astrocytes. Thus, here we wanted to test the hypothesis that LBN has lasting effects on the morphology of astrocytes in the mPFC and OFC. We include females and compare their effects to males, which is an advance over prior studies that have only examined the effects of early life manipulations on astrocytes in males. Another advantage of our approach is that we used a viral labeling strategy with astrocyte-targeted, membrane-tethered Lck-GFP, which unlike GFAP, fully labels the astrocytic processes, so we can obtain a better morphometric picture of stress-induced changes (Testen, Kim et al. 2020). In addition to characterizing astrocyte morphology, we also performed bulk RNA sequencing (RNAseq) on OFC tissue from adult male and female rats exposed to LBN and control manipulations to begin to delineate potential mechanisms by which LBN can alter astrocytes.

## 2.0 Methods

### 2.1 Subjects and the limited bedding and nesting (LBN) manipulation

Male and female Long Evans rats (Charles River Laboratories, Wilmington, MA) were purchased and then bred in-house to prevent additional stress resulting from shipping pregnant dams (Sachs & Lumia, 1981). All dams were primiparous. The day that pups were born was considered PND 0. On PND 2, litters were culled to 10 pups (5 males and 5 females when possible) and litters were randomly assigned to either control housing conditions or LBN housing conditions. Both housing conditions utilized 20×20×40 cm cages with filter tops (non-ventilated). Subjects were placed within LBN housing from PND 2-9, during which a metal grate was placed across the cage floor to prevent access to bedding, there was only one paper towel for nesting (3×10 cm), and no enrichment items were included within their cage (Eck, Ardekani et al. 2019). Subjects within the control condition were placed within standard housing, including ample corncob bedding (about 2 cm deep), two cotton nestlets (5×5 cm each), and one enrichment tube. Throughout, rats were given *ad libitum* access to food and water. Rats were also kept on a 12h reverse light/dark cycle (with lights turning off at 11 am). Weaning occurred at PND21, at which time offspring were pair-housed with same-condition and same-sex partners. Pair-housed rats were moved to a separate colony room on a 12h reverse light/dark cycle (with lights turning off at 9 am). A maximum of 2 rats per sex per litter were utilized for the study. All experiments were approved by and in accordance with guidelines implemented by Temple University’s Institutional Animal Care and Use Committee and the National Institutes of Health guidelines.

### 2.2 Stereotaxic surgery

On the day of the surgery, rats were deeply anesthetized with isoflurane vapor and placed in a stereotaxic instrument to inject the AAV-GfaABC1D-Lck-GFP virus, which selectively labels astrocytes in their entirety, following general protocols outlined in Testen et al., 2020 (Testen, Kim et al. 2020). Unilateral injections of the medial (m) OFC (+4.2 AP, + 0.6 ML, –4.3 DV (measured from the surface of the skull) relative to bregma) and mPFC (+2.9 AP and – 0.6 ML relative to bregma, –4.5 DV measured from the surface of the skull) were performed. Virus was microinjected using an infusion pump (Pump 11 Elite, Harvard Apparatus, Holliston, MA) and 28-gauge injection cannulas (Plastics One, Roanoke, VA). The virus was infused at a rate of 0.1 μl per minute, for a total of 1 μl per injection site. Virus was allowed to diffuse for 15 minutes before injection cannulas were slowly removed over the course of 1 minute. Subjects received an injection of meloxicam (s.c.) in order to mitigate pain at the time of surgery and one day later. Additionally, postoperative care included weighing subjects and inspecting surgical incisions, treating with triple antibiotic ointment if necessary. A timeline for the experimental design in is **Figure 1A**.

**Figure 1.**
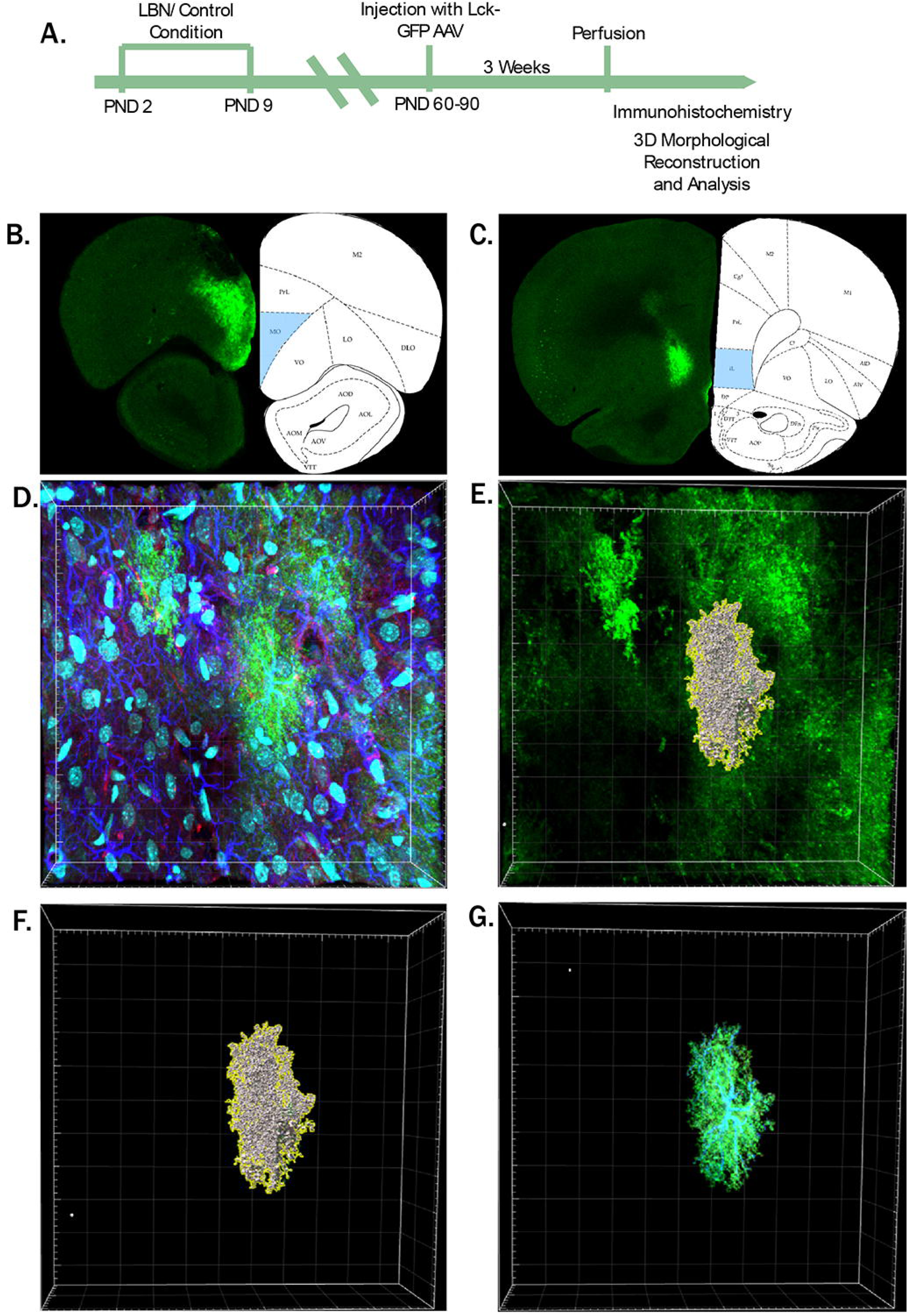
Experimental timeline, representative images of mOFC and mPFC injection sites, and astrocyte reconstruction. (A) Timeline of experimental procedures. (B) mOFC injection sites. (C) mPFC injection sites. (D) Representative image of full confocal image, with AAV-GfaABC1D-Lck-GFP (green), GFAP (dark blue), PSD-95 (red), and DAPI (cyan) staining present. (E) Toggling all other channels off, a “surface” is built around the AAV-GfaABC1D-Lck-GFP signal. This surface serves as an overlay of GFP signaling from the isolated astrocyte, with parameters carefully adjusted to only select valid signal (i.e., no background signal included). (F) Once the surface is built, the 3-D representation of the astrocyte remains. This model is then utilized to analyze morphological and colocalization parameters. (G) “Masked” GFP and GFAP signal. Masked channels can be built, in which all fluorescent signal within the surface is retained, allowing for visualization of the isolated astrocyte.

### 2.3 Tissue collection for morphometric analysis

Following a three-week incubation period to ensure sufficient viral expression (Testen, Kim et al. 2020), brain tissue was collected. Subjects were anesthetized and transcardially perfused with saline heparin followed by 4% paraformaldehyde. Brains were then removed and fixed with 4% paraformaldehyde for 2 h before transferring to a 30% sucrose solution. Brains were stored in the 30% sucrose solution at 2°C for at least 48 h prior to sectioning.

Prior to sectioning, brains were set within a cryostat (Leica CM1950, Wetzlar, Germany) and left to equilibrate to the temperature for at least 45 minutes. Brains were then sliced at 100 µm. Slices containing the mOFC and mPFC were collected and stored within wells containing sufficient cryoprotectant (50% 1× PBS/glycerol) to fully cover the slices. Wells containing samples were wrapped in aluminum foil and stored at –20°C until use for immunohistochemistry.

### 2.4 Immunohistochemistry (IHC)

Prior to beginning IHC, slices were checked for sufficient GFP fluorescence within the regions of interest. To do so, slices near the region of interest were mounted on slides and coverslipped with 1× PBS. Slices were briefly checked using the 20× objective on a fluorescent microscope (Leica DM5500B, Wetzlar, Germany). Once sufficient fluorescence and localization within the region of interest was confirmed (**Fig.1 B,C**), the coverslip was removed and slices were returned to new well plates to be used for immunohistochemistry following the previously published protocol (Testen, Kim et al. 2020). In order to ensure penetrance of antibodies into tissue, only one slice was placed in each well. To minimize the loss of GFP fluorescence, well plates were covered in between each step.

For IHC, slices were first washed 3 × 5 minutes in 1× PBST (0.2% triton x-100) at room temperature on a laboratory shaker. Slices were then blocked in wells containing 5% normal goat serum (Sigma-Aldrich Cat. #G9023) (in 1× PBST (2% triton x-100) for 1 h at room temperature on a laboratory shaker. Slices were then transferred to new wells containing blocking solution with primary antibodies mouse anti PSD-95 (Thermo Fisher Scientific Cat. #MA1-045) at 1:250; rabbit anti-GFAP (Agilent Cat. #Z0334) at 1:500) for 72 h at 4°C on a laboratory shaker. Slices were flipped halfway through the incubation period to ensure optimal penetrance of the antibodies. Following this incubation period, slices were then washed 3 × 5 minutes in 1× PBST (0.2% triton x-100) at room temperature on a laboratory shaker. Slices were then again transferred to new wells containing the blocking solution plus secondary antibodies (goat anti-mouse Alexa Fluor 594 (Thermo Fisher Scientific Cat. #A-11032) at 1:1000; goat anti-rabbit Alexa Fluor 647 (Thermo Fisher Scientific Cat. #A-21245) at 1:1000) for 72 h at 4°C on a laboratory shaker. Again, slices were flipped halfway through the incubation period to ensure optimal penetrance of the antibodies. Following this incubation period, slices were then washed 3 × 10 minutes in 1× PBST (0.2% triton x-100) and 1 × 10 minutes in 1× PBS at room temperature on a laboratory shaker. Slices were then stored in aluminum foil-wrapped wells in cryoprotectant (50% 1x PBS/glycerol) at –20°C until use for imaging. Representative images of immunohistochemical staining are shown in **Supplemental Figure 1**.

### 2.5 Astrocyte imaging

To mitigate any alterations in astrocytic morphology due to drying of brain tissue, slices were mounted and coverslipped immediately prior to imaging. Slices were mounted on Superfrost Plus slides (Fisherbrand, Cat. #22-034-979), and 1-2 drops of a DAPI stain (ACDBio Cat. # 320858) were placed on the slice. DAPI was tapped off the slide after waiting 30 seconds. Slices were then coverslipped with Fluoromount (Thermo Fisher Scientific, Cat. # 00-4958-02).

To analyze astrocytes, slices were then imaged using an Olympus confocal microscope (FV3000, Tokyo, Japan) using 405, 488, 561, and 640 nm diode lasers with a 60× oil-immersed objective and Fluoview software (FV31S-SW, Tokyo, Japan). Frame size was set to 1024 × 1024 pixels and z-step size to 1 µm. Care was taken to only image astrocytes entirely located within the regions of interest (mOFC and mPFC). If astrocytes were located (even partially) within other regions, or appeared to have been cut during sectioning, they were not considered for imaging. If injections missed the regions of interest, slices were not considered for imaging.

When imaging, it was noted that for mPFC infusions, the majority of Lck-GFP expression fell within the infralimbic (IL) region of the mPFC. Because of this, imaging and analysis was restricted to the IL alone to ensure consistency. During image acquisition, researchers remained blinded as to sex and condition in order to ensure unbiased processing.

### 2.6 Morphometric astrocyte analysis

Images were directly imported into the Imaris software (v. 9.9.0., Bitplane, Zürich, Switzerland) with the goal of building a 3-dimensional model of individual astrocytes. Following protocols outlined in Testen et al., 2020, individual astrocytes were isolated, with seven astrocytes analyzed per subject. Surfaces were built upon individual cells by manually adjusting thresholding levels to select Lck-GFP fluorescence from these cells. Care was taken to include Lck-GFP signal from the individual cells alone, excluding any background fluorescence or fluorescence from nearby astrocytes. Using the 3-dimensional model generated from this surface, morphological parameters of interest (namely surface area and volume) were extracted and recorded (**Fig. 1 D-G**).

Following recording of these morphological parameters, masked channels of Lck-GFP and Alexa 594 signal were built. These masked channels consist of signal localized within the 3-dimensional surface alone. Using the masked channels, colocalization analysis was performed between the Lck-GFP and Alexa 594 signal (representing PSD-95 signal), thereby allowing for an analysis of relative astrocyte proximity to postsynaptic excitatory neurons. To accomplish this, the masked Lck-GFP channel was set as the region of interest (ROI) and the masked Alexa 594 channel was set as Channel A (the channel of interest). The threshold for the PSD-95 channel was manually adjusted, with care being taken to remove background noise, but retain PSD-95 puncta. In order to accomplish this, intensity values were averaged for random positive puncta throughout the z-stack. This averaged value was then manually set as the threshold for Alexa 594 signal for the colocalization analysis. After thresholding values were set, the percentage of the ROI colocalized with the Alexa 594 channel was recorded. Importantly, as this parameter is expressed as a percentage of total ROI, it incorporates a control measure for overall size of the astrocyte. During imaging analysis, files were encrypted so that researchers remained blinded as to sex and condition in order to ensure unbiased processing.

### 2.7 Statistical analysis for astrocytes

Collected data were imported into SPSS (v. 28.0.1.0 (142), IBM, Armonk, NY) and were unblinded at this point. Independent variables were sex and condition (control vs. LBN), and dependent variables were surface area, volume, and synaptic colocalization in the area of the astrocyte covered by PSD-95. Outliers for each dependent variable were removed if they fell outside of the standard 1.5 × interquartile region range. Cleaned data were then imported into R (v. 2022.02.1, R Core Team, Vienna, Austria) for further statistical analysis. First, an independent samples t-test was performed to see if there were significant differences between cortical layers in any of the morphological or PSD-95 related parameters, as previous literature demonstrated that there may be significant heterogeneity in morphometric properties of astrocytes between layers (Lanjakornsiripan et al., 2018). All the mOFC astrocytes analyzed were located within cortical layer V. mPFC astrocytes were located in layers V and VI, but no significant differences were found between these two layers, so analysis proceeded without taking layers into account. Within R, nested ANOVA analysis was performed for all groups for assessment of morphological parameters, synaptic co-localization, and expression of PSD-95 puncta. All figures were generated within Prism (v. 9.0.0. (121), GraphPad Software, San Diego, California).

### 2.8 Bulk RNA sequencing of the OFC

RNA sequencing was conducted to assess gene expression changes across sexes and two conditions from OFC: Controls and LBN. Quality control (QC) was conducted using Fastqc software (version 0.11.8) (Andrews 2010). The Trimmomatic (version 0.39) software (Bolger, Lohse et al. 2014) was then applied to remove non-paired reads and identified adaptors. RNAseq reads were mapped on rn6 genome assembly using the Hisat2 (Sirén, Välimäki et al. 2014) as the alignment step, assembled by StringTie (Pertea, Pertea et al. 2015). The differentially expressed genes (DEGs) were selected by applying the Deseq2 library (Love, Huber et al. 2014) in R (version 4.3.1) (R Core Team 2020). The conditions to define DEGs were a 50% change in gene expression (|log 2 Fold change| > 0.58) of genes and an adjusted p-value < 0.1. Also, for downstream analysis and visualization of the RNA-seq analysis output, including, but not limited to, drawing heatmaps and Venn diagrams, R statistical software (version 4.3.1) was used as previously described (Sahraeian, Mohiyuddin et al. 2017, Sanchez, Bavley et al. 2021). HOMER (the software for motif discovery and next-generation sequencing analysis) was used to find the DEGs’ transcription factors (Heinz, Benner et al. 2010). The applied criteria was − 2000 to + 1000 bp of the transcriptional start site and a length of 8–12 bp for TFs.

Gene set enrichment analysis (GSEA) is a method to recognize sets of genes or proteins over-represented in a large group of genes or proteins and may associate with specific phenotypes (e.g., LBN) (Subramanian, Tamayo et al. 2005). GSE analysis was done using the g: Profiler website service as previously described (Reimand, Isserlin et al. 2019), using DEGs from groups separately. Pathways (from KEGG) and gene ontology terms (biological process) by adjusted p-value < 0.1 were selected for further analysis. To compare samples’ gene expression agnostically, the Rank–Rank Hypergeometric Overlap (RRHO) analysis was used. This algorithm steps within two gene lists ranked by the p-value of differential expression observed in two experiments and estimate the number of overlapping genes. Subsequently, a heatmap shows the strength and correlation pattern between the two expression profiles (Plaisier, Taschereau et al. 2010). This analysis used the RRHO2 library in R (Cahill, Huo et al. 2018) to compare the extent of overlap in gene expression patterns in males and females.

To identify specific sub-cell types that contribute to changes observed in a bulk RNA-seq differential gene expression experiment, the LRcell deconvolution package was used(Sharma, Ma et al. 2023). This method used sets of cell marker genes captured from single-cell RNA-sequencing as indicators for various cell types in the tissue of interest. Next, for each cell type, it applied Logistic Regression on the complete set of genes with differential expression p-values to calculate a cell-type significance p-value. Finally, these p-values were compared to predict which one(s) were likely to be responsible for the differential gene expression pattern observed in the bulk RNA-seq experiments. Given our focus on astrocyte mechanisms, we used this approach to identify DEGs in our data set astrocyte-enriched markers.

## 3.0 Results

### 3.1 LBN causes astrocyte enlargement in the mOFC

Brian sections collected from LBN, and control animals were subjected to morphometric astrocyte analyses (**Fig. 1**, also see methods for details). Five subjects were analyzed for each control group, and 4 subjects were analyzed for each LBN group, with 7 cells analyzed per subject. For the examination of mOFC astrocytic surface area, group n’s were as follows, denoted as n subjects/n astrocytes: F/CTL n= 5/35, M/CTL n= 5/34, F/LBN n=4/27, M/LBN n= 4/26, leaving 1 M/CTL, 1 F/LBN, and 2 M/LBN cells that were excluded as outliers. For the examination of mOFC astrocytic volume, group n’s were as follows: F/CTL n= 5/32, M/CTL n= 5/34, F/LBN n= 4/25, M/LBN n= 4/24, leaving 3 F/CTL, 1 M/CTL, 3 F/LBN, and 4 M/LBN cells that were excluded as outliers. This analysis demonstrated that astrocytes within the mOFC of both male and female subjects that had experienced LBN exhibited significantly increased surface area (F(1, 14) = 4.65, p = .049) and volume (F(1, 14) = 8.878, p = .0099) (**Fig. 2 A,B**).

**Figure 2.**
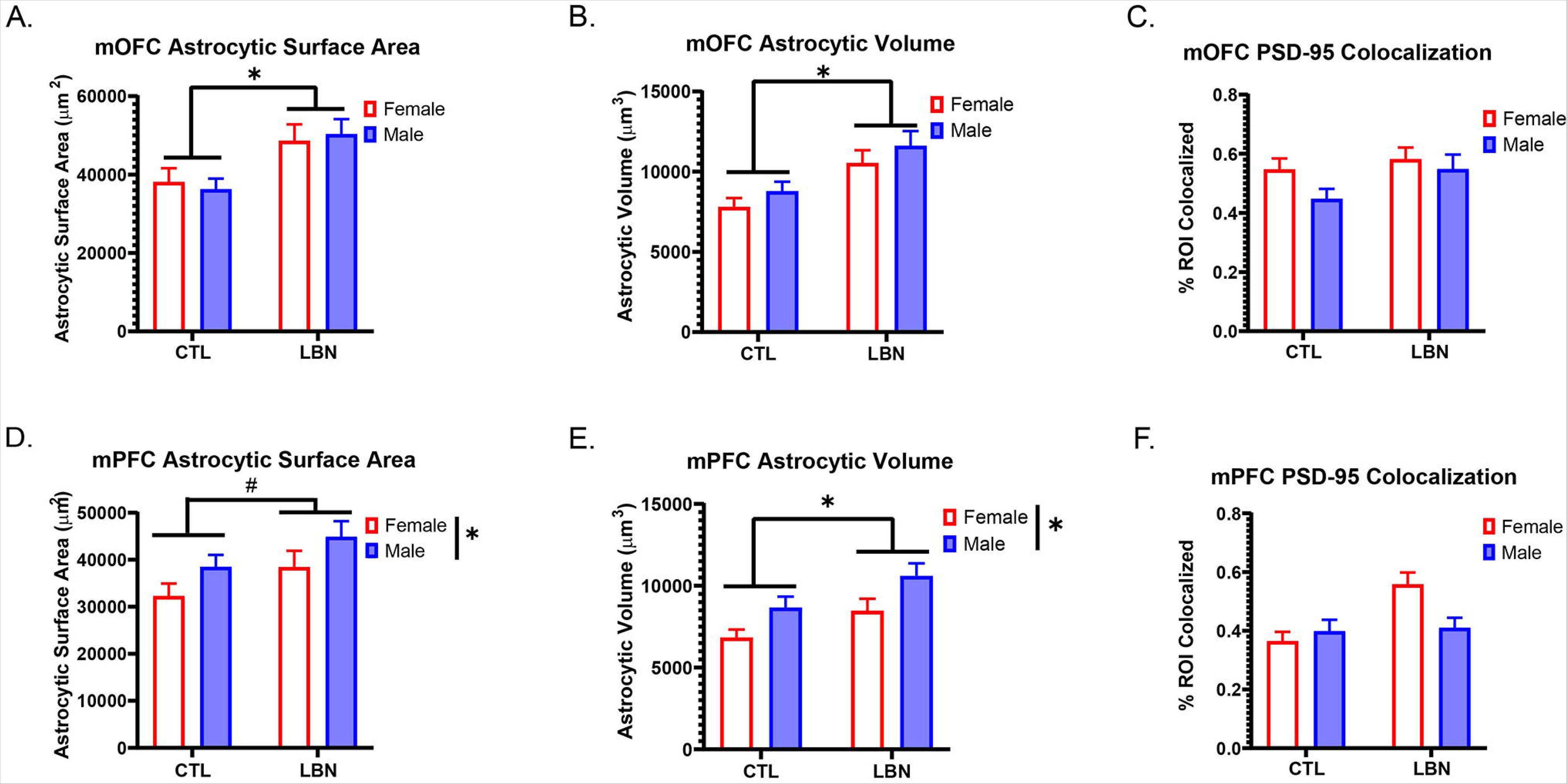
mOFC and mPFC astrocytic morphology and PSD95 colocalization. (A) LBN increases associated with increased astrocytic surface area in the mOFC in both male and female subjects. (B). The experience of LBN is associated with increased astrocytic mOFC volume in both male and female subjects. (C) There are no significant differences in colocalization in PSD-95 colocalization with the isolated GFP signal. (D) Quantification of mPFC astrocytic surface area. There is a main effect of sex, with males having larger astrocytic surface area than females. Additionally, there is a trend towards a main effect of condition, with astrocytic surface area being larger in subjects that have experienced LBN. (E) The experience of LBN is associated with increased mPFC astrocytic volume in both male and female subjects. Additionally, there is a significant main effect of sex, with astrocytic volume being larger in males. (F) There are no significant differences in colocalization between PSD-95 and the isolated GFP signal in any of the groups. Asterisks represent p<.05. # indicate p<.07. Error bars represent SEMs.

For the PSD-95 colocalization analysis, group n’s were as follows: F/CTL n= 5/34, M/CTL n= 5/32, F/LBN n= 4/27, M/LBN n= 4/26, leaving 1 F/CTL, 3 M/CTL, 1 F/LBN, and 1 M/LBN cells that were excluded as outliers. This colocalization analysis between GFP– and PSD-95 fluorescence demonstrated that there were no significant differences in PSD-95 colocalization between any of the groups in terms of condition (F(1,14) = 1.26, p = .280), sex (F(1,14) = 1.378, p = .0.26), or an interaction between the two (F(1,14) = 0.33, p = .575) (**Fig, 2C**).

### 3.2 LBN causes astrocyte enlargement in the mPFC

Four subjects were analyzed for each group, with 7 cells analyzed per subject. For the examination of mPFC astrocytic surface area, group n’s were as follows, denoted as n subjects/n astrocytes: F/CTL n= 4/26, M/CTL n= 4/26, F/LBN n= 4/26, M/LBN n= 4/25, leaving 2 F/CTL, 2 M/CTL, 2 F/LBN, and 3 M/LBN cells that were excluded as outliers. For the examination of mPFC astrocytic volume, group n’s were as follows: F/CTL n= 4/26, M/CTL n= 4/27, F/LBN n= 4/27, M/LBN n= 4/25, leaving 2 F/CTL, 1 M/CTL, 1 F/LBN, and 3 M/LBN cells that were excluded as outliers. When analyzed, a trend towards increased astrocytic surface area in male and female LBN subjects was observed, but this trend did not reach statistical significance, (F(1,12) = 4.20, p = .063) (**Fig. 2D**). Additionally, astrocytes within mPFC of both male and female subjects that had experienced LBN exhibited significantly increased astrocytic volume, (F(1,12) = 6.36, p = .027) (**Fig. 2E**). Furthermore, a trend for a main effect of sex was present, with astrocytic surface area (F(1,12) = 4.41, p = .058) and there was a significant main effect of sex for volume (F(1,12) = 8.542, p = .0123) with overall larger astrocytes in males as compared to females (**Fig. 2 D,E**). This baseline sex difference in astrocyte size within the mPFC is consistent with previous work (Bollinger, Salinas et al. 2019), which indicates that male Sprague-Dawley rats have larger astrocytes in the mPFC than females.

For the PSD-95 colocalization analysis, group n’s were as follows: F/CTL n= 4/26, M/CTL n= 4/25, F/LBN n= 4/26, M/LBN n= 4/26, leaving 2 F/CTL, 3 M/CTL, 2 F/LBN, and 2 M/LBN cells that were excluded as outliers. Colocalization analysis between GFP– and PSD-95 fluorescence demonstrated that there were no significant differences in PSD-95 colocalization between any of the groups in terms of condition (F(1, 12) = 3.60, p = .082), sex (F(1,12) = 0.935, p = .353), or an interaction between the two (F(1,12) = 2.62, p = .131) (**Fig. 2F**).

### 3.3 LBN alters the OFC transcriptome differently in male and female rats

We conducted bulk RNA-sequencing on punches from OFC from adult behaviorally naïve rats (M/CTL, n = 5; F/CTL, n = 5; M/LBN, n = 5; F/LBN, n = 5) to look for molecular signatures in the OFC that could account for the observed change in astrocyte morphology. Although LBN resulted in larger cortical astrocytes in males and females, we analyzed these data within sex (i.e., comparing LBN v. control for males and comparing LBN v. control for females). Sex-specific analysis is warranted because there can be latent sex differences, where the physiological change is similar across sex, but the underlying mechanisms are different (De Vries 2004, Oberlander and Woolley 2016).

We first used rank-rank hypergeometric overlap (RRHO) analysis to compare overall gene expression patterns after LBN in males and females. RRHO facilitates an agnostic comparison of gene expression patterns because it includes gene changes that fail to reach statistical significance. Remarkably, there are no hotspots for shared LBN-induced upregulated and downregulated genes in males and females in the OFC (**Fig. 3A**; bottom left and top right quadrants). Instead, LBN induced unique signatures in males and females with the strongest hotspot for genes that are upregulated in males and downregulated in females (**Fig. 3A**; hotspot bottom right quadrant). We next used a combination of fold change (|log 2 Fold change| > 0.58) and FDR corrected p-values (*p*<.1) to define differentially expressed genes (DEGs). LBN resulted in 179 DEGs in males and 156 DEGs in females (**Fig. 3B, Supplemental Table 1**).

**Figure 3.**
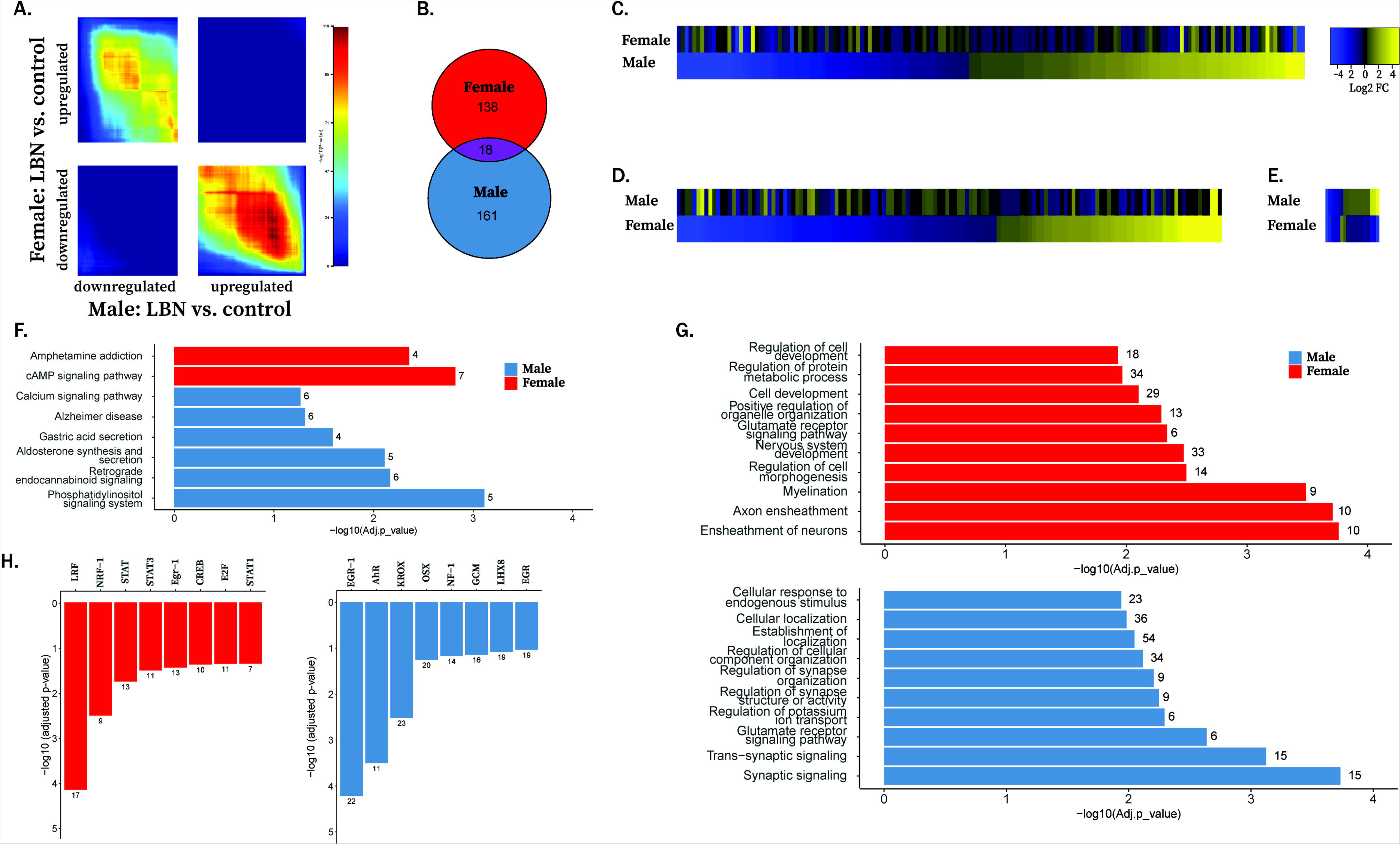
LBN produces changes in gene expression in the OFC of males and females. (A) Rank-rank hypergeometric overlap computing threshold-free comparison of gene expression. Hotspots represent the overlap between the impact of LBN on gene expression in females and males with hotter colors representing more overlap. The upper right quadrant includes co– upregulated genes and co-down regulated genes are represented in the lower left quadrant. (B-E) LBN produced 156 DEGs in females and 179 DEGs in males with 18 or those overlapping between sexes. (C) Heatmaps sorted by fold change highlighting the differences between females (C) and males (D) or overlapping DEGs (E). (F) Biological Process enrichment terms in males and females following LBN. (G) KEGG enrichment terms identified in males and females following LBN. (H) Potential master regulators of LBN-induced changes in gene expression, predicted using HOMER.

Heatmaps sorted by DEG fold change revealed that LBN induced different patterns of up-regulated and down-regulated genes across sex: in males, 90 of identified DEGs were up-regulated following LBN, while in females, only 59 of DEGs were up-regulated (**Fig. 3C, D**). Of the 18 overlapping DEGs, only 10 (7 downregulated and 3 upregulated) changed in the same direction in both sexes (**Fig. 3E**). Thus, even though LBN had a similar effect on astrocyte morphology in the OFC, LBN caused sex-divergent transcriptional changes in this region.

To determine biological processes altered by LBN, we subjected DEGs to Kyoto Encyclopedia of Genes and Genomes (KEGG) and Biology Process (BP) ontology analysis. We focused on pathways involved in neurotransmission, signaling, development, metabolism, neuropsychiatric/neurodegenerative disease, and the immune system as these pathways were most implicated in brain function (Walker, Zhou et al. 2022). We included relevant KEGG pathways that reached significance (adj *p-value* <.10), and these pathways mostly differed between males and females (**Fig. 3F, Supplemental Table 2**). Long-term potentiation (LTP) and dopaminergic synapse were significant in both males and females, but different sets of DEGs were regulated by LBN within these pathways: LTP (males: *Prkca*, *Itpr1*, *Gnaq*; females: *Camk2d*, *Grin1*, *Grin2b*) and dopamine synapse (males: *Drd2, Prkca*, *Itpr1*, *Gnaq*; females: *Camk2d*, *Grin2b*, *Akt2*, *Ppp2r5b*, *Gng13).* For BP, we only included the top 10 relevant enrichment terms per analysis and saw that most terms only reached significance in one sex (**Fig. 3G**). An exception was the glutamate receptor signaling pathway that appeared in males and females. Yet again, the genes enriched by LBN in this pathway were largely sex-specific (males: *Daglb*, *Ncstn*, *Gnaq*, *Mef2c*, *Shank3*, *Park7*; females: *Arc*, *Grin1*, *Grin2b*, *Neto1*, *Grid1*, *Park7*), except for *Park7* which was upregulated by LBN in both sexes.

Hypergeometric Optimization of Motif Enrichment (HOMER) analysis of transcription factor binding sites was performed to identify potential master regulators affected by LBN that could account for the broad effects on transcription in the OFC. HOMER analysis found largely different transcription factor binding sites for the DEGs altered by LBN in males and females (**Fig. 3H**). However, the transcription factor Egr1 (aka, Zif268) was affected by LBN across sex, and Egr transcription factors regulate astrocyte proliferation (Mayer, RoLssler et al. 2009). **Figure 4** illustrates LBN-induced sex differences in aspects of the glutamate receptor signaling pathway in the OFC.

**Figure 4.**
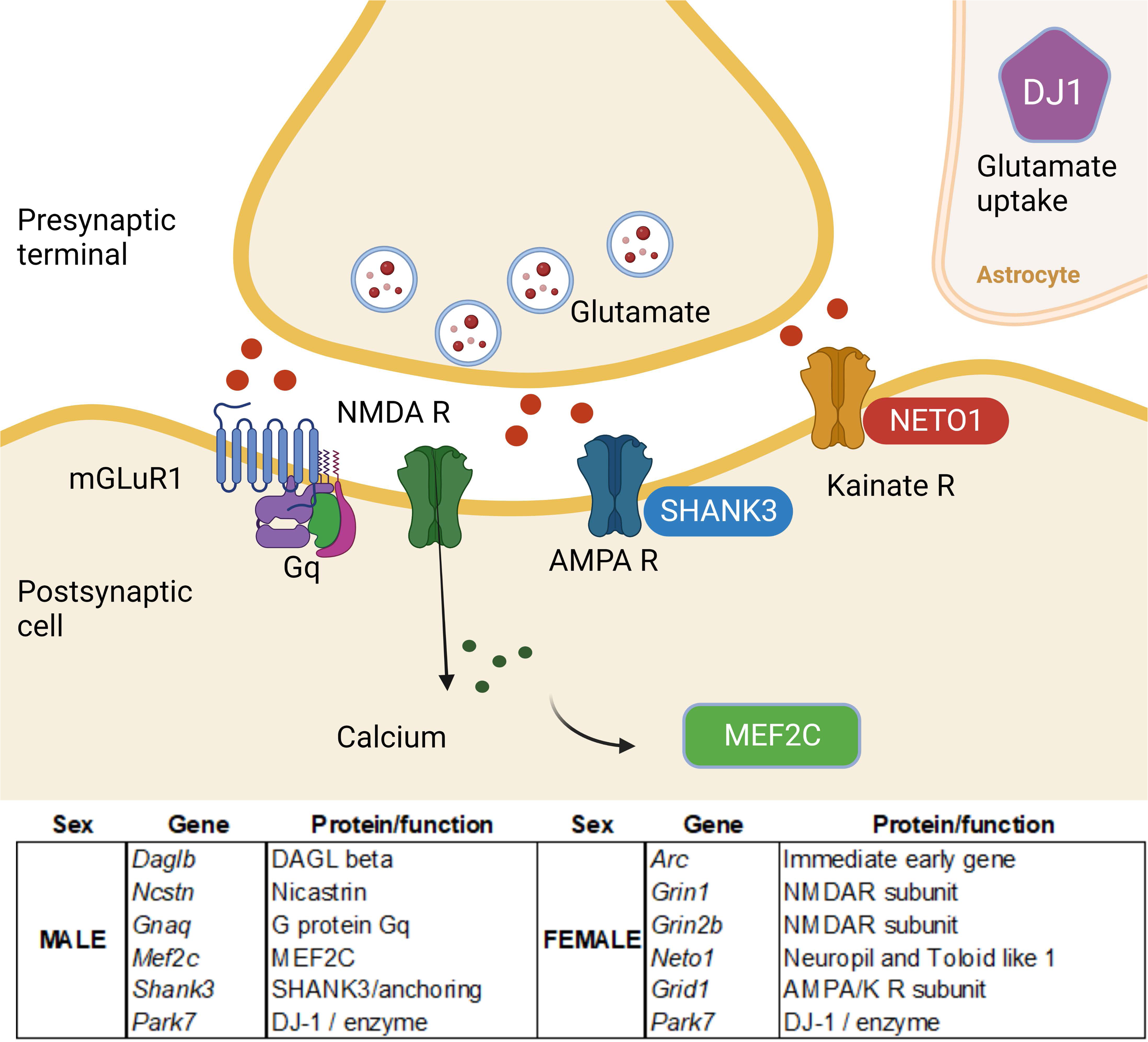
LBN affects signaling molecules involved in glutamatergic signaling in the OFC of males and females. Top of figure shows the location and function of proteins encoded by DEGs that produced enrichment in glutamate signaling enrichment term in males and females. The table lists the genes in males and females and their corresponding proteins and function in neurons. Created in part with Biorender.com

Results of LRcell deconvolution package showed that in LBN significantly enriched genes in astrocytes in the OFC of males and females (FDR corrected p= .004 and.02, respectively). The genes found primarily in astrocytes that were altered by LBN in males are, *Fermt2*, *Gpm6a*, and *Tmem229a*, and for females are, *Fxyd1*, *Car2*, *Hhatl*, *Akt2*, *Wscd1*, *Tmem229a*, and *Gdf10*. Of note, *Gpm6a* encodes for GPM6A (neuronal membrane glycoprotein M6A), which promotes synaptogenesis and spine formation and is downregulated by prenatal stress in male rats (Monteleone, Adrover et al. 2014, León, Aparicio et al. 2021). Unlike prenatal stress, which is largely associate with later vulnerability, LBN promotes later stress resilience. Here we found that *Gpm6a* is upregulated by LBN in males, an effect that may help astrocytes maintain synapses as they get larger.

Here we found that LBN causes astrocyte enlargement in the OFC. Yet it is difficult to tell from morphology alone whether this hypertrophy represents healthy plasticity or reactive astrocytes, which respond to pathological stimuli such as injury or disease. Moreover, there is no prototypical reactive astrocyte, thus there is not a single marker that can define them. Rather, there are a number of putative markers of reactive astrocytes (Escartin, Galea et al. 2021). Given the number of transcripts determined by RNAseq, we leveraged our data to assess whether LBN altered the transcription of a suite of reactive astrocytic markers, ranging from cytoskeletal elements to transporters (based on (Escartin, Galea et al. 2021). **Table 1** shows the LBN-induced log fold change (positive number indicates an upregulation by LBN) along with the FDR corrected p-values in males and females for reactive astrocyte markers. In some cases, p-values could not be calculated due to low coverage of a transcript in a particular gene. All the p-values were *p*>.99, so did not approach significance. This pattern suggests that LBN-induced astrocyte hypertrophy is not indicative of reactive astrocytes.

## 4.0 Discussion

Here we found that brief exposure to early resource scarcity using the LBN manipulation causes lasting changes in astrocyte morphology, increasing the volume and surface area of astrocytes in mOFC and mPFC in adult male and female rats. Despite the fact that morphological changes were similar across sex, there was sex-specific regulation of the OFC transcriptome by LBN. One possibility is that the few shared transcriptional changes that occur across sex could be the key genes that alter morphology. Another possibility is that latent sex differences are at play. For example, LBN alters different genes in the glutamate signaling pathway in males versus females, yet the net effect, altered glutamate signaling, may be similar across sex. Glutamate signaling mediates astrocyte morphogenesis (Morel, Higashimori et al. 2014, Torres-Ceja and Olsen 2022), so this LBN-induced change in signaling could result in similar morphological changes in male and female astrocytes, but via sex-specific mechanisms.

The consequences of cortical astrocyte enlargement due to LBN remain to be determined. LBN and control rats had similar amounts of PSD-95 colocalization with astrocytes. This result suggests that LBN does not cause a compensatory reduction in excitatory synapses to counteract the increase in the size of the astrocytes. Instead, the PSD-95 density remains similar, but with the increased size of the astrocytes in the LBN group, the overall effect would likely be enhanced regulation of cortical glutamatergic signaling by astrocytes in the LBN group. Of course, astrocytes have many other functions, so LBN also likely alters glutamate turnover, neuronal metabolism, and other homeostatic functions in cortex via increased astrocytic morphology. Whether these changes are neuroprotective, and perhaps account for the stress-resilient cortically mediated behavior phenotypes we see following LBN (e.g., reduced impulsivity and risk taking), will need to be investigated in future studies. However, we found no evidence that LBN significantly increased putative markers of reactive astrocytes, suggesting that this change in morphology is not pathological.

### 4.1 LBN causes lasting cortical astrocytic morphology

Here we demonstrate that LBN causes astrocytes in the mOFC and mPFC to get larger. Although this is the first study to assess LBN effects in cortical astrocytes, there have been a couple reports of LBN altering astrocytes in different brain regions (Gunn, Cunningham et al. 2013, Abbink, Kotah et al. 2020). In the mouse hippocampus, there are developmental changes in GFAP coverage following LBN (Abbink, Kotah et al. 2020). Specifically, LBN increases GFAP coverage in the stratum lacunosum-moleculare layer of the hippocampus at PND9, but this effect is transient and not observed at young adult and middle-aged timepoints (1, 4, and 6 months). However, at 10 months, GFAP coverage throughout the hippocampus is reduced in LBN versus controls (Abbink, Kotah et al. 2020). This change in astrocyte coverage is not accompanied by any changes in hippocampal astrocyte morphology (Abbink, Kotah et al. 2020). However, in the neonatal mouse hypothalamus, LBN thins GFAP-immunoreactive arbors (Gunn, Cunningham et al. 2013). This effect is associated with impaired astrocytic glutamate uptake in corticotropin releasing factor (CRF)-expressing dorsalmedial neurons (Gunn, Cunningham et al. 2013). Both studies assessed morphology with GFAP so likely missed important morphological features. These prior studies also assessed the effect of LBN on astrocytes at earlier timepoints and, for the Abbink et al. study, across development. In contrast, we only looked at changes in young adult rats, so our interpretation is limited to this age. It is possible if we looked at neonatal or aged timepoints, then we would observe different, perhaps dynamic, changes in cortical astrocyte morphology. Given the crucial function of astrocytes to brain development and age-related disorders such as Alzheimer’s disease (Garwood, Ratcliffe et al. 2017, Farhy-Tselnicker and Allen 2018, Habib, McCabe et al. 2020), future studies should assess how LBN causes morphological changes of astrocytes in many brain regions across development.

Relative to the sparse data on early life adversity, there is more work on chronic adult stress and cortical astrocytes, which typically demonstrates stress-induced reductions in GFAP (Banasr, Chowdhury et al. 2010, Sántha, Veszelka et al. 2016, Shilpa, Bhagya et al. 2017).

However, chronic restraint stress had no effect on astrocyte number but reduced astrocyte morphology as assessed with GFAP (Tynan, Beynon et al. 2013). Similarly, here our transcriptomic analysis found no significant change in GFAP expression in the OFC, yet LBN altered astrocyte morphology in this region. Thus, morphological changes may be more sensitive to stress than overall astrocyte number.

A difference between our early resource scarcity findings and adult chronic stress findings is that we found cortical astrocyte hypertrophy while adult chronic stress causes reduced astrocyte number or atrophy. This distinction likely maps onto stress vulnerable versus resilient phenotypes. The aforementioned adult chronic stressors typically cause anxiety– and depression-like behaviors, as well as cognitive deficits (Banasr, Chowdhury et al. 2010, Sántha, Veszelka et al. 2016, Shilpa, Bhagya et al. 2017). However, our LBN manipulation reduces impulsivity, risky decision making, and opioid self-administration, while increasing social self-administration, with these phenotypes typically more pronounced in male rats (Ordoñes Sanchez 2021, Ordoñes Sanchez, Bavley et al. 2021, Williams, Flowers et al. 2022). Therefore, we view our brief early stressor as promoting resilience, consistent with the phenomena of stress inoculation, where early adversity that is not overwhelming can lead to better stress coping later in life (Masten 2001, Lyons, Parker et al. 2010). Given that the astrocyte changes observed here are not indicative of reactive astrocytes, our interpretation is that the LBN-induced astrocyte growth helps promote these resilient phenotypes by facilitating cortical glutamate transmission, turnover, and overall metabolism. Indeed, in LBN animals, PSD95 coverage of astrocytes were maintained at control levels. That LBN animals maintain PSD95 cover while astrocyte get larger suggests that overall brief early scarcity increases astrocyte interactions with excitatory synapses, which may facilitate glutamate signaling. In futures studies, we hope to directly test this idea and changes in other functions mediated by cortical astrocytes following LBN.

### 4.2 Potential mechanisms underlying astrocyte enlargement following LBN

To begin to explore mechanisms by which early scarcity causes lasting changes in astrocyte morphology, we performed RNAseq on the OFC to identify transcriptional changes may drive this plasticity. A striking outcome of these analyses was the widespread sex-specific changes in gene transcription following LBN. This contrasts with the similar effects of LBN on astrocyte enlargement in the OFC across sex. One possibility is that the few shared gene changes across sex are the key drivers of changes in astrocyte morphology. For example, LBN upregulated *Park7* in the OFC of male and female rats. *Park7* encodes for the DJ-1 protein, and deletion of *Park7* causes familial Parkinson’s disease (Bonifati, Rizzu et al. 2003). DJ-1 is highly expressed in astrocytes (Bandopadhyay, Kingsbury et al. 2004, Bader, Ran Zhu et al. 2005), and its overexpression in this cell type augments neuroprotective mechanisms against oxidative stress (Mullett and Hinkle 2009, Mullett, Di Maio et al. 2013). In contrast, knocking out DJ-1 impairs astrogliosis, as DJ-1 knock out mice show less astrocytic hypertrophy around a brain injury than wild type animals (Choi, Eun et al. 2018). Although currently unknown, it is possible that increasing DJ-1 levels increases astrocytic morphology. If so, the increase in *Park7* expression following LBN may drive cortical astrocyte enlargement in males and females, and also would be expected to promote resilience to oxidative stress.

Another possibility is that the increased in cortical astrocyte morphology following LBN in males and females occurs via latent sex differences. As noted, astrocytes regulate glutamate signaling, but in turn glutamate signaling mediates astrocyte morphogenesis (Morel, Higashimori et al. 2014, Torres-Ceja and Olsen 2022). Specifically, during development synaptically released glutamate acts as signal to promote the maturation of cortical astrocytes (Morel, Higashimori et al. 2014). Here we found that LBN altered glutamate signaling genes. However, the transcripts within the glutamate pathway that were affected by LBN differed in males versus females.

Future studies need to employ more direct measures of glutamate signaling following LBN in cortex, but it is possible that glutamate signaling is increased across sex, but the mechanism that led to the increase are sex-specific representing a latent sex difference.

### 4.3 Astrocyte vulnerability to early life manipulations

It may seem surprising that a brief exposure to scarcity causes changes lasting in cortical astrocyte morphology that are observed in adulthood. However, we implement LBN at a critical time of astrocyte proliferation, cortical colonization, and maturation (Farhy-Tselnicker and Allen 2018, Clavreul, Abdeladim et al. 2019). The work on the cortical development and maturation has primarily occurred in mouse models, however the developmental timeline for rat is likely similar. During late embryonic development of the mouse (embryonic days 11-18) is when neurogenesis and migration occur, processes that precede astrogenesis (Farhy-Tselnicker and Allen 2018, Clavreul, Abdeladim et al. 2019). While some astrocyte proliferation happens during late embryonic development, the first postnatal week of life is when astrogenesis peaks. From PND 0-7 pre– and postnatal astrocyte progenitors colonize the cortex then proliferate locally (Clavreul, Abdeladim et al. 2019). Later in the postnatal period (PND 7-21), there is a maturation phase where, instead of proliferating, astrocytes gain morphological complexity and establish a continuous cortical astrocyte network (Clavreul, Abdeladim et al. 2019). After that, the proliferative capacity of astrocytes drops, except when reactive astrocytes proliferate and migrant to a site of injury (Catalani, Sabbatini et al. 2002, Sofroniew and Vinters 2010, Clavreul, Abdeladim et al. 2019, Pavlou, Grandbarbe et al. 2019). Given we implement LBN from PND 2-9, changes in neurochemistry due to scarcity and/or altered parental care could directly impact cortical astrogenesis and maturation. How LBN causes lasting effects on astrocytes remains to be determine, but their proliferation, differentiation, and function is regulated via transcriptional and epigenetic processes (Pavlou, Grandbarbe et al. 2019). Future studies will assess genetic and epigenetic effects of LBN at the time of LBN exposure and compare alterations in astrocytes across the development.

## 4.4 Conclusions

Most of the field has focused on how LBN alters neurons or microglia. However, given that the timing of this model coincides with a sensitive period for astrocyte proliferation and maturation, astrocytes may be a cell type particularly sensitive to the effects of LBN. Here we demonstrate that in cortex, LBN causes astrocyte enlargement in adulthood. While the mechanisms underlying this change are not fully delineated, we identified LBN-induced alterations in transcripts in astrocyte genes (e.g., *Park7*) and glutamate signaling pathways that could contribute to the morphology changes in the LBN group. Given the role of astrocytes in a range of functions from regulating glutamate signaling to glucose metabolism, these changes in morphology likely have a big impact on cortical brain function, which we plan to assess in future studies. The present work underscores that the field needs to better consider the impact of the early environment on astrocytes to fully understand the mechanisms by which early life events cause lasting changes in plasticity and behavior.

## 5.0 Acknowledgements

We would like to thank Molly Dupuis for her technical assistance. This research includes calculations carried out on Temple University’s HPC resources and thus was supported in part by the National Science Foundation through major research instrumentation grant number 1625061 and by the US Army Research Laboratory under contract number W911NF-16-2-0189.

The research is supported by grants from the National Science Foundation, IOS-1929829 to DAB, and the National Institute of Health, MH129020 and DA049837 to DAB, DA056534 and DA055846 to DAB and MEW. Some of the figures were created using Biorender.com

## Notes

### Competing Interest Statement

The authors have declared no competing interest.

