## Supplemental Figure and Tables for "Early resource scarcity causes cortical astrocyte enlargement and sex-specific changes in the orbitofrontal cortex transcriptome in adult rats"

### *Supplemental Figure A.* Representative immunohistochemical staining.

A.

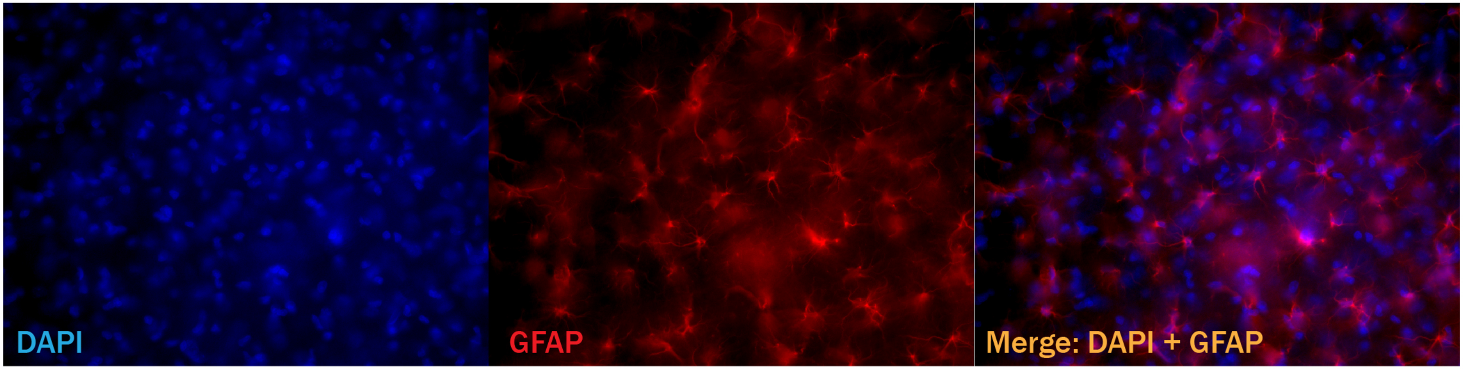


B.

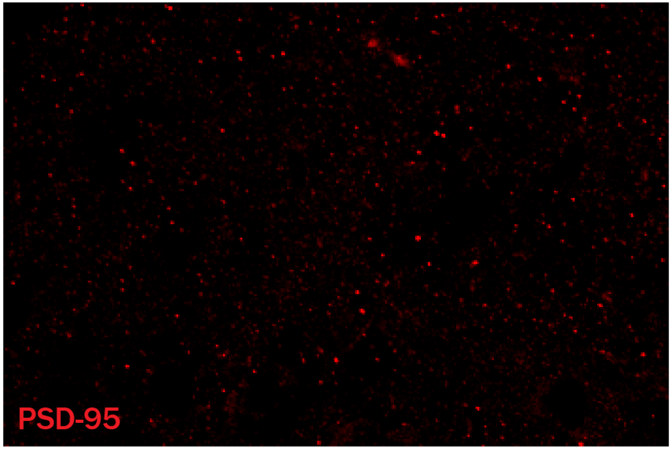


Supplemental Figure A. Representative immunohistochemical staining. (A) Immunohistochemical staining at 40x. From left to right, DAPI staining, GFAP staining, and merged image are shown. (B) Representative image of PSD-95 staining at 60x, allowing for visualization of individual PSD-95 puncta.

### Supplemental Table 1. Differentially Expressed Genes (DEGs)

| Male DEGs | | | Female DEGs | | |
|---|---|---|---|---|---|
| Gene symbol | log2FoldChange | Adj.pvalue | Gene symbol | log2FoldChange | Adj.pvalue |
| Acot7 | 22.4967 | 0.00000 | Sept8 | 22.0838 | 0.00000 |
| AABR07007916.1 | -4.1111 | 0.00001 | Pnkd | -6.5024 | 0.00003 |
| Kcnab2 | -3.4672 | 0.00025 | Rps21 | 2.0669 | 0.00014 |
| Tnk2 | -2.1612 | 0.00043 | AC109958.1 | -3.9748 | 0.00050 |
| Mef2c | -5.8409 | 0.00054 | Necab3 | -4.5112 | 0.00175 |
| Fam219a | -3.8681 | 0.00077 | Plekho1 | -2.2305 | 0.00259 |
| Mfsd13a | 5.0910 | 0.00104 | Eif3c | -1.5680 | 0.00323 |
| Lonrf2 | 3.7632 | 0.00245 | Ss18l2 | -3.1482 | 0.00328 |
| Abcf2 | 4.8002 | 0.00252 | Dap3 | 2.6775 | 0.00361 |
| Rplp0 | 1.7136 | 0.00288 | AABR07059891.1 | -6.3404 | 0.00395 |
| Epha5 | -2.4450 | 0.00321 | Rps6kc1 | 3.0120 | 0.00413 |
| Pou3f2 | -3.9127 | 0.00339 | Rpl3 | -1.6269 | 0.00419 |
| Wnk2 | 6.4076 | 0.00351 | Arnt2 | -5.7376 | 0.00598 |
| Klhl26 | -4.7562 | 0.00377 | Fem1a | 5.9852 | 0.00612 |
| Psenen | -2.4829 | 0.00513 | Jak1 | -2.6207 | 0.00640 |
| AC134224.3 | -6.0972 | 0.00605 | Fam13b | -5.8952 | 0.00678 |
| Plcd3 | 3.1781 | 0.00629 | Phf12 | -3.4203 | 0.00868 |
| AC134224.1 | 3.7487 | 0.00730 | Megf9 | -5.5826 | 0.01023 |
| Arhgef2 | 1.6110 | 0.00843 | Eif4e | -3.9941 | 0.01072 |
| Plekho1 | 3.5739 | 0.00963 | Rnf44 | -2.7612 | 0.01136 |
| Camk2n1 | -7.8015 | 0.00998 | Plekhg5 | 3.2445 | 0.01279 |
| Pnkd | -5.6308 | 0.01144 | Rap1gap | 3.1189 | 0.01298 |
| Rangap1 | 3.1718 | 0.01204 | Agpat4 | 2.7251 | 0.01357 |
| Pdcd11 | -3.0829 | 0.01213 | Nt5dc3 | 3.7305 | 0.01386 |
| Rpl18 | 1.9978 | 0.01238 | Arc | 0.8009 | 0.01399 |
| Mpnd | -3.0402 | 0.01280 | Slc25a42 | -2.2303 | 0.01639 |
| Ndufs7 | 2.5331 | 0.01320 | Zpr1 | -2.5847 | 0.01671 |
| Igsf1 | 3.6150 | 0.01397 | Smurf1 | 2.0076 | 0.01823 |
| Kcnq5 | -5.2816 | 0.01505 | Mir1188 | 3.9317 | 0.01827 |
| Rps6ka4 | -1.2416 | 0.01659 | RGD1308428 | -2.8646 | 0.01877 |
| RT1-S3 | -1.7553 | 0.01705 | Hddc2 | -2.7272 | 0.01942 |
| Cops7a | -3.1611 | 0.01720 | Gabbr1 | -3.7971 | 0.01961 |
| Mgst3 | -2.7563 | 0.01749 | Eral1 | -4.9777 | 0.02191 |
| Peg3 | 7.0164 | 0.01797 | Park7 | 0.8820 | 0.02197 |
| AABR07064061.1 | -2.9297 | 0.01832 | Tango2 | -3.9240 | 0.02233 |
| Chst10 | 3.4050 | 0.01882 | AABR07071891.2 | 1.8594 | 0.02284 |
| Ash1l | -4.4030 | 0.02080 | Dync1h1 | 5.7349 | 0.02307 |
| Gpr137 | -6.8740 | 0.02081 | Cirbp | -1.2427 | 0.02344 |
| Mink1 | -5.0069 | 0.02111 | Zfp870 | -2.9957 | 0.02468 |
| Pde2a | 1.7519 | 0.02267 | Grin2b | 2.5836 | 0.02482 |
| Ly6h | 0.9072 | 0.02316 | Lingo3 | 1.4934 | 0.02519 |
| Def8 | 2.8302 | 0.02341 | Gid8 | 2.5033 | 0.02523 |
| Dlg3 | -5.3325 | 0.02366 | Noc2l | -3.0457 | 0.02543 |
| Retreg3 | 1.5936 | 0.02492 | Bbs9 | -2.6600 | 0.02582 |
| AABR07044362.6 | -4.8646 | 0.02494 | Foxj3 | 1.8184 | 0.02584 |
| Ccdc82 | 3.3824 | 0.02522 | Tmem229a | -4.8707 | 0.02664 |
| RF01268 | 1.4962 | 0.02561 | LOC100911576 | -2.4424 | 0.02706 |

| | | | | | |
|---|---|---|---|---|---|
| Fermt2 | -4.8391 | 0.02584 | Tmem125 | -1.6167 | 0.02838 |
| Prepl | -1.9078 | 0.02788 | Hspe1 | 5.2385 | 0.02879 |
| Nxph3 | 0.9283 | 0.02839 | Piga | -3.6989 | 0.02889 |
| Kdelc1 | 2.8927 | 0.02884 | Mypop | -4.7052 | 0.03139 |
| Trak2 | 2.6327 | 0.02941 | Tcp1 | -3.5769 | 0.03142 |
| Dtnb | -1.4752 | 0.03330 | Cpne9 | 2.2118 | 0.03203 |
| LOC100911196 | 3.0106 | 0.03367 | Grin1 | 6.4129 | 0.03215 |
| Pex5l | -6.2538 | 0.03413 | Josd2 | -4.1285 | 0.03267 |
| Daglb | 1.8772 | 0.03430 | Neto1 | -1.7777 | 0.03295 |
| Fam166a | 4.6802 | 0.03473 | Actg1 | 1.7044 | 0.03343 |
| Mtg1 | 4.6034 | 0.03489 | Myo5a | -1.6673 | 0.03386 |
| Snx3 | -2.5507 | 0.03490 | Sik2 | -2.4873 | 0.03430 |
| LOC100360573 | 3.1517 | 0.03529 | AABR07001382.1 | 0.7981 | 0.03472 |
| Nampt | -2.3586 | 0.03644 | Map1lc3b | -3.9221 | 0.03503 |
| Park7 | 0.8146 | 0.03709 | Atp13a1 | -2.3625 | 0.03627 |
| Zc3h4 | 2.5494 | 0.03725 | Cadm2 | -1.3866 | 0.03826 |
| Zc3hc1 | -4.5384 | 0.03727 | Mapt | 3.8199 | 0.03965 |
| Slc25a42 | -1.8255 | 0.03770 | Wscd1 | -1.1851 | 0.04001 |
| Sparc | -0.8821 | 0.03782 | Stx7 | -4.2271 | 0.04047 |
| Rprm | 0.6797 | 0.03888 | Timm29 | 4.9101 | 0.04082 |
| LOC100911951 | 2.2633 | 0.03965 | Nr4a1 | 0.7064 | 0.04175 |
| Gyg1 | 3.5016 | 0.04016 | Arpc2 | -3.4367 | 0.04208 |
| Golgb1 | 2.5510 | 0.04069 | Fxyd1 | -2.7338 | 0.04303 |
| Rbm42 | 1.0942 | 0.04321 | Pam | 1.7696 | 0.04418 |
| LOC108348112 | -1.4190 | 0.04422 | Lasp1 | -1.2301 | 0.04418 |
| Spock2 | -2.0046 | 0.04492 | Rbfox3 | 1.5021 | 0.04499 |
| LOC100910750 | 2.7699 | 0.04513 | Nxph3 | -0.9336 | 0.04511 |
| Gkap1 | -3.0801 | 0.04863 | Slc37a3 | -4.0593 | 0.04512 |
| Ddx39a | 2.2665 | 0.04960 | Snx3 | -2.4025 | 0.04597 |
| Fis1 | -3.1458 | 0.05096 | Tnip1 | -3.6892 | 0.04687 |
| Dgkb | -2.8090 | 0.05126 | Fis1 | -3.3810 | 0.04706 |
| Strada | 3.0807 | 0.05247 | Pcnx1 | 1.8885 | 0.04829 |
| Creld1 | -0.8623 | 0.05257 | Zfp330 | -0.9715 | 0.04850 |
| Cpsf4 | 2.9163 | 0.05260 | Wdr45 | -3.7041 | 0.04891 |
| Myh11 | -3.0862 | 0.05368 | Pitpnb | -2.4050 | 0.04918 |
| Atxn2 | 2.5794 | 0.05409 | Ifnar1 | 2.3911 | 0.05006 |
| Mtmr4 | 1.3866 | 0.05557 | Arl6ip5 | -3.8127 | 0.05033 |
| Strip2 | -2.0510 | 0.05559 | Mpp3 | -3.1768 | 0.05059 |
| Rbfox3 | -1.1837 | 0.05705 | Fbxl3 | 4.4493 | 0.05131 |
| Atp5pb | -1.3653 | 0.05758 | Mobp | -0.9172 | 0.05233 |
| Slain1 | 2.7963 | 0.05949 | Ehmt2 | -3.6200 | 0.05329 |
| Usp31 | -2.3033 | 0.05975 | Ap4s1 | -3.3369 | 0.05332 |
| Scoc | -1.8800 | 0.06090 | Akt2 | 2.9370 | 0.05734 |
| Mboat7 | 1.4853 | 0.06098 | Cldn11 | -1.0240 | 0.05737 |
| Slco3a1 | 2.2033 | 0.06115 | Eef2k | -2.7685 | 0.05764 |
| Pnpla8 | -2.3814 | 0.06123 | Lemd3 | -1.9258 | 0.05882 |
| Gnaq | 0.9654 | 0.06220 | Mal | -0.9548 | 0.06122 |
| Nsmf | 2.2983 | 0.06279 | Dgcr6 | 1.2077 | 0.06230 |
| Igsf8 | 0.6098 | 0.06355 | Ganab | 2.1827 | 0.06343 |
| AABR07054368.1 | 5.4402 | 0.06414 | Nat14 | -1.7270 | 0.06367 |
| Ncstn | 2.2723 | 0.06590 | Cttn | 3.8740 | 0.06374 |

| | | | | | |
|---|---|---|---|---|---|
| Adora1 | 1.1769 | 0.06617 | Aars | 1.7130 | 0.06434 |
| Gpm6a | 1.6365 | 0.06627 | Cnp | -0.8155 | 0.06447 |
| Dmac2 | -2.0612 | 0.06665 | AABR07073186.1 | 5.5040 | 0.06530 |
| Gabra5 | -1.1240 | 0.06715 | Hhatl | 3.2941 | 0.06679 |
| Rpl32 | 0.8664 | 0.06762 | Pdp1 | 0.7663 | 0.06704 |
| Scn4b | -0.9071 | 0.06795 | Wnk2 | -5.4684 | 0.06738 |
| Uck1 | 3.3674 | 0.06831 | Tspan14 | -1.0864 | 0.06756 |
| Ncbp1 | -1.8730 | 0.06953 | C2cd4b | -2.4178 | 0.06769 |
| Eno1 | 0.9981 | 0.06992 | Tspan2 | -0.9167 | 0.06833 |
| Itsn1 | 3.3458 | 0.07007 | Camk2d | 2.9830 | 0.06941 |
| Spata2L | -1.5044 | 0.07007 | RGD1561149 | 1.9156 | 0.07135 |
| Rps6kc1 | -2.9800 | 0.07034 | Fktn | -2.0796 | 0.07278 |
| Dpy19l3 | -3.5788 | 0.07059 | Fxn | -1.9206 | 0.07292 |
| Prrt2 | -1.4261 | 0.07070 | Mrpl34 | 5.3268 | 0.07473 |
| Drd2 | -2.3352 | 0.07073 | Mbp | -0.8801 | 0.07520 |
| Dync1i2 | 3.2685 | 0.07109 | Amotl1 | -1.6710 | 0.07565 |
| Fem1a | 5.3361 | 0.07142 | Dcun1d2 | 1.3619 | 0.07581 |
| Pam | 1.8520 | 0.07285 | Cdc42ep2 | -2.2803 | 0.07662 |
| Lemd3 | -1.6601 | 0.07350 | Uqcrb | 0.8953 | 0.07673 |
| Shc1 | -2.4937 | 0.07380 | Cnpy2 | -1.7856 | 0.07743 |
| Ezh1 | -1.4868 | 0.07423 | Nap1l4 | 5.2773 | 0.07808 |
| Pdyn | -1.1315 | 0.07427 | Rbbp4 | 3.3658 | 0.07823 |
| Mta1 | -2.7307 | 0.07444 | AABR07018038.1 | -2.3224 | 0.07868 |
| Gle1 | -2.6636 | 0.07505 | Grid1 | -2.6512 | 0.07918 |
| Mafg | -2.0919 | 0.07620 | Mog | -0.9802 | 0.07969 |
| Ssx2ip | 5.2485 | 0.07657 | Rasd2 | 0.7239 | 0.08025 |
| Snph | 5.2313 | 0.07743 | Vps53 | 5.2336 | 0.08070 |
| Mbp | 0.7291 | 0.07748 | Tmem88b | -1.0440 | 0.08124 |
| Rnf165 | -2.7332 | 0.07772 | Evi2a | -0.9082 | 0.08191 |
| Dnajc9 | -2.3392 | 0.07811 | Adora2a | 2.5433 | 0.08232 |
| Ntn3 | 3.1910 | 0.07898 | Bcas1 | -0.7286 | 0.08241 |
| Mx2 | 2.3606 | 0.07911 | Ppm1e | -0.5863 | 0.08251 |
| Coq8a | 2.5317 | 0.07959 | Ppp2r5b | -0.6207 | 0.08300 |
| Tmem229a | 2.7622 | 0.07992 | Mir212 | 1.0070 | 0.08313 |
| AABR07017250.1 | 2.0698 | 0.08108 | Abcg1 | -1.1595 | 0.08475 |
| Lmf1 | -3.8677 | 0.08269 | Klk6 | -1.2129 | 0.08533 |
| Dpp6 | -0.6471 | 0.08353 | Hivep2 | -1.0067 | 0.08758 |
| Vars | 1.5103 | 0.08377 | Gng13 | -1.2861 | 0.08799 |
| Mtmr12 | 3.4265 | 0.08392 | Dlg1 | 0.6652 | 0.08867 |
| Arhgap20 | -1.1318 | 0.08459 | Dmac2 | -1.8704 | 0.08937 |
| Sec63 | 1.7868 | 0.08495 | Psme4 | 1.6576 | 0.08970 |
| Miga2 | 2.5820 | 0.08497 | Grn | 1.8283 | 0.09058 |
| Caskin2 | 1.9387 | 0.08498 | Fam189b | 1.9234 | 0.09110 |
| Tnpo3 | -1.3932 | 0.08544 | Fa2h | -0.9677 | 0.09111 |
| Cct6a | -1.0519 | 0.08569 | Tp53bp2 | 1.5898 | 0.09152 |
| Tmcc1 | 2.2783 | 0.08586 | Smc6 | -1.2205 | 0.09226 |
| Tspan17 | 0.8571 | 0.08628 | Gdf10 | -2.6984 | 0.09247 |
| Cnot1 | 2.5003 | 0.08672 | Rpl26 | -0.8550 | 0.09359 |
| LOC108348161 | 2.0243 | 0.08768 | Egr3 | 1.1190 | 0.09444 |
| Sirpa | 1.2357 | 0.08798 | Ptpn4 | 2.2271 | 0.09476 |
| Ccn2 | 1.1152 | 0.08883 | Ppp1r14a | -0.7842 | 0.09552 |

| | | |
|---|---|---|
| Tpm1 | -2.0510 | 0.08902 |
| Zpr1 | -1.9866 | 0.08928 |
| Shank3 | -0.9054 | 0.08945 |
| Tppp3 | 0.6736 | 0.08952 |
| Actr2 | -1.1076 | 0.08963 |
| Ilvbl | -2.2318 | 0.08990 |
| Ninj2 | 1.4104 | 0.09132 |
| Fcho1 | -3.0747 | 0.09134 |
| Prkca | -0.8875 | 0.09174 |
| Itpr1 | 1.4592 | 0.09208 |
| Emc6 | -4.9893 | 0.09294 |
| Jup | -0.9222 | 0.09344 |
| Bhmt | 0.9293 | 0.09359 |
| mrpl9 | -4.1168 | 0.09400 |
| Gtpbp2 | -2.9513 | 0.09488 |
| LOC100362965 | 4.4975 | 0.09573 |
| Mpp3 | 2.3928 | 0.09578 |
| RF00003 | 2.3895 | 0.09591 |
| P2rx7 | 2.3885 | 0.09596 |
| Abcb6 | -1.7417 | 0.09659 |
| Glt8d1 | -2.9274 | 0.09666 |
| Cdk11b | -1.8294 | 0.09675 |
| Flrt1 | 1.1545 | 0.09697 |
| Pip4k2b | 1.2671 | 0.09721 |
| Tln2 | -1.9644 | 0.09794 |
| Cyb561 | -1.3640 | 0.09843 |
| Cckbr | -1.9487 | 0.09932 |
| Zdhhc22 | 0.9340 | 0.09948 |
| Tle5 | -1.3083 | 0.09973 |
| Apbb1 | -0.6630 | 0.09583 |
| Ap1ar | 1.4959 | 0.09685 |
| Tor1aip1 | -2.5117 | 0.09731 |
| AABR07037489.1 | -2.2776 | 0.09755 |
| AABR07021988.1 | -2.2801 | 0.09759 |
| Car2 | -0.6512 | 0.09881 |
| Lpgat1 | -2.6821 | 0.09947 |

### Supplemental Table 2. Enrichment

| Sex | Sorce | Pathway/Term | Adj p-value | No of genes | Gene names |
|---|---|---|---|---|---|
| Male | KEGG | Phosphatidylinositol signaling system | 0.0008 | 5 | PRKCA ITPR1 MTMR4 PLCD3 DGKB |
| | | Retrograde endocannabinoid signaling | 0.0068 | 6 | PRKCA ITPR1 GABRA5 DAGLB NDUFS7 GNAQ |
| | | Aldosterone synthesis and secretion | 0.0078 | 5 | PRKCA ITPR1 DAGLB GNAQ PDE2A |
| | | Alzheimer disease | 0.0489 | 6 | ATP5PB ITPR1 NCSTN PSENEN NDUFS7 GNAQ |
| | | Calcium signaling pathway | 0.0542 | 6 | PRKCA ITPR1 CCKBR PLCD3 P2RX7 GNAQ |
| | | Gap junction | 0.0598 | 4 | DRD2 PRKCA ITPR1 GNAQ |
| | | Morphine addiction | 0.0691 | 4 | PRKCA ADORA1 GABRA5 PDE2A |
| | | CGMP-PKG signaling pathway | 0.0695 | 5 | ITPR1 ADORA1 MEF2C GNAQ PDE2A |
| | | Glutamatergic synapse | 0.0814 | 4 | PRKCA ITPR1 SHANK3 GNAQ |
| | | Cholinergic synapse | 0.0826 | 4 | PRKCA ITPR1 KCNQ5 GNAQ |
| | | Long-term depression | 0.0847 | 3 | PRKCA ITPR1 GNAQ |
| | | Parathyroid hormone synthesis, secretion and action | 0.0887 | 4 | PRKCA ITPR1 MEF2C GNAQ |
| | | Long-term potentiation | 0.0903 | 3 | PRKCA ITPR1 GNAQ |
| | | Dopaminergic synapse | 0.0947 | 4 | DRD2 PRKCA ITPR1 GNAQ |
| | | Thyroid hormone synthesis | 0.0979 | 3 | PRKCA ITPR1 GNAQ |
| | | Renin secretion | 0.0999 | 3 | ITPR1 ADORA1 GNAQ |
| | BP | Synaptic signaling | 0.0002 | 15 | P2RX7 DRD2 DAGLB DLG3 GABRA5 CAMK2N1 ADORA1 NCSTN PREPL NSMF PNKD PRRT2 DGKB MEF2C SHANK3 |
| | | Trans-synaptic signaling | 0.0008 | 15 | DRD2 DAGLB DLG3 GABRA5 CAMK2N1 ADORA1 P2RX7 NCSTN PREPL NSMF PNKD PRRT2 DGKB MEF2C SHANK3 |
| | | Glutamate receptor signaling pathway | 0.0023 | 6 | DAGLB NCSTN GNAQ MEF2C SHANK3 PARK7 |
| | | Regulation of potassium ion transport | 0.0051 | 6 | WNK2 ADORA1 DRD2 KCNAB2 LOC100911951 DPP6 |
| | | Regulation of synapse structure or activity | 0.0057 | 9 | GPM6A SPARC ITSN1 DRD2 FLRT1 DGKB MEF2C SHANK3 ACTR2 |
| | | Regulation of synapse organization | 0.0062 | 9 | GPM6A SPARC ITSN1 DRD2 FLRT1 DGKB MEF2C SHANK3 ACTR2 |
| | | Regulation of cellular component organization | 0.0077 | 34 | GPM6A SPARC ATXN2 SIRPA GLE1 TPPP3 P2RX7 ITSN1 DRD2 TPM1 CCT6A FIS1 TNK2 EPHA5 ACTR2 POU3F2 CNOT1 PIP4K2B CCN2 CDK11B ARHGEF2 RETREG3 RPS6KA4 PLEKHO1 FLRT1 NCBP1 PRRT2 DGKB MEF2C SHANK3 TRAK2 MBP PAM DEF8 |
| | | Establishment of localization | 0.0090 | 54 | P2RX7 SEC63 SIRPA GABRA5 TRAK2 ZDHHC22 KCNQ5 GLE1 ATP5PB WNK2 ZPR1 TNPO3 NCBP1 NDUFS7 FCHO1 ADORA1 DRD2 KCNAB2 LOC100911951 LMF1 CCT6A ATXN2 FIS1 TNK2 ITSN1 EPHA5 DDX39A ACTR2 ITPR1 PREPL STRADA GPM6A TMCC1 PNKD JUP SSX2IP MBP CCKBR PARK7 ABCB6 ARHGEF2 SLC25A42 MTMR12 SCN4B PRRT2 DPP6 RANGAP1 SLCO3A1 MEF2C PNPLA8 SHANK3 RPLP0 PAM CCN2 |
| | | Cellular localization | 0.0104 | 36 | P2RX7 SEC63 DLG3 TRAK2 ZDHHC22 GLE1 ZPR1 TNPO3 NCBP1 LMF1 CCT6A FIS1 ITSN1 EPHA5 DDX39A ACTR2 ITPR1 PREPL DRD2 STRADA FERMT2 TMCC1 PNKD JUP SSX2IP PARK7 TLN2 ARHGEF2 PRRT2 GOLGB1 RANGAP1 MEF2C SHANK3 ADORA1 KCNAB2 DEF8 |
| | | Cellular response to endogenous stimulus | 0.0115 | 23 | LY6H LEMD3 RNF165 FIS1 RANGAP1 EPHA5 MTMR4 NSMF FERMT2 CNOT1 CCN2 JUP PARK7 ZPR1 GKAP1 PDE2A ARHGEF2 FLRT1 MEF2C RPLP0 ACTR2 NAMPT RPL32 |
| Female | KEGG | cAMP signaling pathway | 0.001509624 | 7 | CAMK2D GRIN1 GRIN2B AKT2 ADORA2A FXYD1 GABBR1 |
| | | Amphetamine addiction | 0.004375633 | 4 | CAMK2D GRIN1 GRIN2B ARC |
| | | Dopaminergic synapse | 0.004959881 | 5 | CAMK2D GRIN2B AKT2 PPP2R5B GNG13 |
| | | Circadian entrainment | 0.006635717 | 4 | CAMK2D GRIN1 GRIN2B GNG13 |
| | | PI3K-Akt signaling pathway | 0.033748766 | 7 | EIF4E AKT2 IFNAR1 PPP2R5B GNG13 NR4A1 JAK1 |
| | | Long-term potentiation | 0.044263799 | 3 | CAMK2D GRIN1 GRIN2B |
| | | Alzheimer disease | 0.083979339 | 5 | GRIN1 GRIN2B MAPT APBB1 UQCRB |
| | BP | Ensheathment of neurons | 0.000174033 | 10 | MAL MBP CLDN11 ZPR1 AKT2 FA2H TSPAN2 DLG1 MYO5A BCAS1 |
| | | Axon ensheathment | 0.000195016 | 10 | MAL MBP CLDN11 ZPR1 AKT2 FA2H TSPAN2 DLG1 MYO5A BCAS1 |
| | | Myelination | 0.000324346 | 9 | MAL MBP ZPR1 AKT2 FA2H TSPAN2 DLG1 MYO5A BCAS1 |
| | | Regulation of cell morphogenesis | 0.003181699 | 14 | CDC42EP2 SS18L2 SMURF1 MAPT GRIN1 ARPC2 PLEKHO1 CPNE9 AP1AR DLG1 ARC EEF2K MBP CTTN |
| | | Nervous system development | 0.003352256 | 33 | RBFOX3 MAL MBP AARS ZPR1 SS18L2 ADORA2A PPP2R5B KLK6 SMURF1 MAPT CLDN11 GRIN1 ARNT2 NECAB3 CNP EGR3 AKT2 FA2H CPNE9 TSPAN2 EHMT2 DLG1 EIF4E MYO5A PAM CTTN ARC RAP1GAP EEF2K APBB1 BCAS1 MOBP |
| | | Glutamate receptor signaling pathway | 0.004595877 | 6 | ARC GRIN1 GRIN2B NETO1 GRID1 PARK7 |
| | | Positive regulation of organelle organization | 0.005128601 | 13 | CDC42EP2 SMURF1 FIS1 MAPT DYNC1H1 TCP1 ARPC2 AKT2 GRN AP1AR PPM1E DLG1 CTTN |
| | | Cell development | 0.007927879 | 29 | ZPR1 SS18L2 PSME4 ADORA2A PPP2R5B KLK6 SMURF1 MAPT GRIN1 ARPC2 CNP AKT2 FA2H GRN CPNE9 TSPAN2 AP1AR EHMT2 ACTG1 EIF4E ARC CAMK2D RAP1GAP EEF2K MBP NECAB3 APBB1 CTTN TCP1 |
| | | Regulation of protein metabolic process | 0.010781193 | 34 | SMURF1 CIRBP AARS DCUN1D2 PPP2R5B NOC2L PPM1E ADORA2A RASD2 FIS1 CNPY2 MAPT ARL6IP5 NR4A1 TNIP1 EEF2K MBP WNK2 PARK7 AKT2 HHATL PPP1R14A GRN PLEKHG5 FKTN DLG1 GDF10 EIF4E PSME4 GRIN1 CAMK2D EHMT2 APBB1 FEM1A |
| | | Regulation of cell development | 0.011684886 | 18 | SS18L2 PPP2R5B KLK6 SMURF1 MAPT GRIN1 ARPC2 GRN CPNE9 AP1AR EIF4E ARC RAP1GAP EEF2K MBP NECAB3 APBB1 CTTN |
